# *mCherry* contains a fluorescent protein isoform that interferes with its reporter function

**DOI:** 10.1101/2021.12.07.471677

**Authors:** Maxime Fages-Lartaud, Lisa Tietze, Florence Elie, Rahmi Lale, Martin Frank Hohmann-Marriott

**Affiliations:** Department of Biotechnology, Norwegian University of Science and Technology, Trondheim, N-7491, Norway; United Scientists CORE (Limited), Dunedin 9016, Aotearoa - New Zealand

## Abstract

Fluorescent proteins are essential reporters in cell biology and molecular biology. Here, we reveal that red-fluorescent proteins possess an alternative translation initiation site that produces a short functional protein isoform. The short isoform creates significant background fluorescence that biases the outcome of expression studies. Our investigation identifies the short protein isoform, traces its origin, and determines the extent of the issue within the family of red fluorescent protein. Our analysis shows that the short isoform defect of the red fluorescent protein family may affect the interpretation of many published studies. Finally, we provide a re-engineered mCherry variant that lacks background expression as an improved tool for imaging and protein expression studies.

## Introduction

The mathematician Gottlob Frege dedicated his life to demonstrate that mathematics were reducible to logic. In June 1901, his ambitions were shattered by a letter from Bertrand Russell that pointed out a contradiction in his fundamental assumption that became famously known as “Russell’s paradox” [1]. Russell’s paradox states he following: Consider *R* the set of all sets that do not contain themselves as members; if *R* is not a member of itself, then by definition, it is a member of itself, and reciprocally.

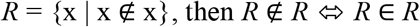

This contradiction left Frege in a state of epistemic paralysis [2], questioning the foundation of his lifelong hypothesis. Interestingly, Russell wrote that Frege responded to the challenge of his main hypothesis with intellectual pleasure and dedication to advance knowledge [1]. We encountered an equivalent of Russell’s paradox in molecular biology, more specifically in relation with the expression of the red fluorescent protein “mCherry”.

The discovery of fluorescent proteins has played a major role in unravelling details of cellular functions [3,4]. Fluorescent proteins display structural similarities, such as a fully amino acid-encoded chromophore present on an α-helix that is tightly packed into an eleven-stranded β-barrel [5,6]. The chromophore is created by the autocatalytic cyclization of an amino acid triade [6–8]. The discovery of DsRed expanded the color range of fluorescent proteins to include red wavelengths [9]. The natural versions of fluorescent proteins have been subject to a multitude of modifications to obtain different colors [6,10–12] and improve their properties such as solubility, maturation, stability, quantum yield, monomeric state or the ability to uptake a fusion partner [13–17]. This diversity of engineered fluorescent proteins have emerged as invaluable tools for molecular biology and cell biology, as they are excellent reporters for gene expression and subcellular protein localization in various biological systems [3,4,15].

In a previous study, we used mCherry as one of the reporters for the development of a universal gene expression method that employs 200 random nucleotides as regulatory sequence to drive expression of a coding sequence [18]. The efficiency of the method is gene- and context-dependent, but it yields usually between 30% and 40% of successful protein expression in *E. coli* [18]. However, when *mCherry* is used as a reporter, we observe fluorescence in about 65% of *E. coli* clones (*Supplemental Figure S1*). Additionally, a large-scale analysis of the transcription start sites (TSS) of these clones showed a non-negligible fraction of leaderless mRNA sequences [18]. Unsettled by these high proportions, our suspicion turned to the reporter sequence itself. We hypothesized that internal Shine-Dalgarno (SD) sequences within the reporter sequence followed by methionine codons just downstream the actual start codon, could result in expression of a shorter yet still functional version of mCherry. This situation reminded us of Russell’s paradox: The fluorescent protein mCherry does not contain itself; however, we suspected that it does (*Supplemental Figure S2*).

A fraction of eukaryotic and prokaryotic genes possess alternative translation initiation sites (ATIS) that lead to the production of different isoforms of a functional protein from a unique mRNA [19–22]. The N-terminal sequence variation between protein isoforms can be the target of post-translational regulation [23], affect protein functionality [24,25] and even direct subcellular localization [26]. The presence of an ATIS in *mCherry* greatly affects its function as a reporter and the outcome of experiments. A shorter version of mCherry contained in itself creates a non-negligible background fluorescence, disturbing the results of gene expression and protein localization studies. For example, a genetic construct encoding a fusion protein composed of a C-terminal mCherry presents a risk of producing an independent, fusion-less mCherry protein, which interferes with protein localization. Likewise, studies using *mCherry* as reporter for gene expression would yield biased results because translation of the short mCherry isoform is included in the reporter gene sequence. As *mCherry* is widely used, the postulated interference may affect many studies, including our own work. However, like Frege, we believe that the advancement of knowledge deserves our full dedication. Therefore, we investigated *mCherry* expression in detail to uncover the source of the issue with the goal of providing solutions that eliminate the defect.

## Results and Discussion

### Identification of the shorter protein isoform of mCherry

Our previous experiments with *mCherry* as a reporter gene [18] lead us to suspect the presence of a shorter protein isoform of mCherry. We first analyzed the sequence of the codon-optimized version of *mCherry* and noticed three methionine residues in a relatively close proximity from the annotated start codon (*Figure 1*). In addition, we recognized an SD-like sequence ranging from -12 to -6 nucleotides upstream of the first internal methionine residue (*Figure 1*). This led us to hypothesize that a short functional isoform of mCherry is produced from one of the three downstream methionine and not from an alternative start codon.

**Figure 1.**
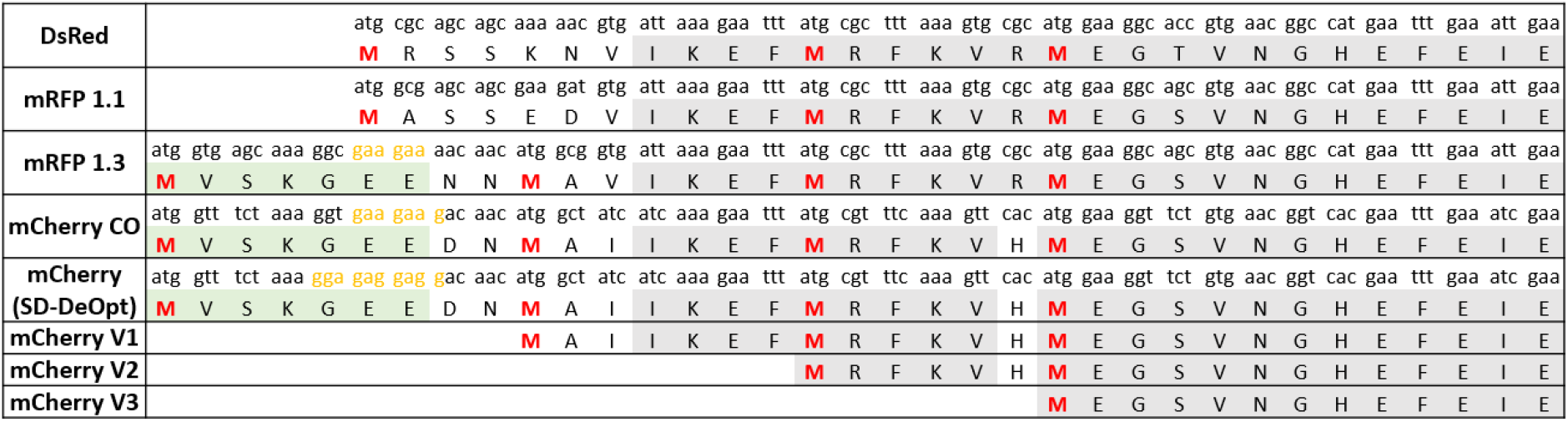
Amino acid sequence alignment of key engineering steps of red fluorescent proteins derived from DsRed. The isoform-causing N-terminal extension occurred in the engineering step from mRFP1.1 to mRFP1.3, due to of the addition of the eGFP fragment (in green) and the linker NNMA. The sequences of the short mCherry versions (V1, V2 and V3) used to determine the shortest functional isoform are also presented. The sequence of the “SD-de-optimized” version of mCherry is shown. Methionine potentially serving as start codons are shown in red. SD-like sequences are colored in yellow. Codon usage of each protein is indicated above each amino acid. Conserved amino acid sequence is shadowed in gray.

In order to assess which of these methionine residue functions as an ATIS that still renders a functional mCherry protein, we designed three versions of mCherry (version V1, V2 and V3), with increasing N-terminal truncations (*Figure 1*). Each version of *mCherry* was expressed in *E. coli* with a constitutive promoter/5’-UTR. The fluorescence measurements of the different mCherry versions are presented in Figure 2. The smallest truncation (version V1) retains 73% of the fluorescence intensity of the original codon optimized version (*mCherry-CO*), while the other versions (V2 and V3) do not show any fluorescence. Additionally, proteomics analysis of the red fluorescent protein mCherry V1 confirmed that the first 10 amino acids were absent (*Supplemental Figure S3*), although the fragment between M10 and M17 could not be detected, due to the short length of the digested fragments. This is conclusive evidence that the *mCherry* gene produces a short functional protein isoform starting at the methionine in position 10 of the amino acid sequence.

**Figure 2.**
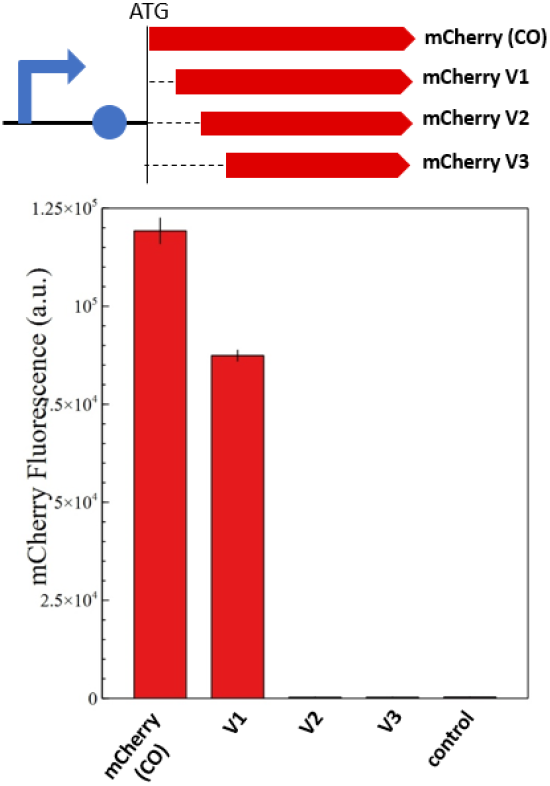
Fluorescence measurements of mCherry and its shorter isoforms (V1, V2 and V3). The genetic map of each constitutively expressed *mCherry* version is shown above the fluorescence quantification graph. A plasmid without *mCherry* is used as negative control. Only *mCherry V1* produces a functional protein isoform that retains 73% of the original fluorescence.

### Phylogenic analysis reveals the origin and the extent of the problem

Since most red fluorescent proteins originate from modifications of DsRed [9], we imagined that the dual-isoform issue of mCherry could affect other members of the red fluorescent protein family. We performed protein sequence alignments to determine the extent of the issue across DsRed-derived fluorescent proteins (*Figure 1*). We were able to trace back the apparition of the second isoform to the engineering of mRFP1.3 (*Figure 3*).

**Figure 3.**
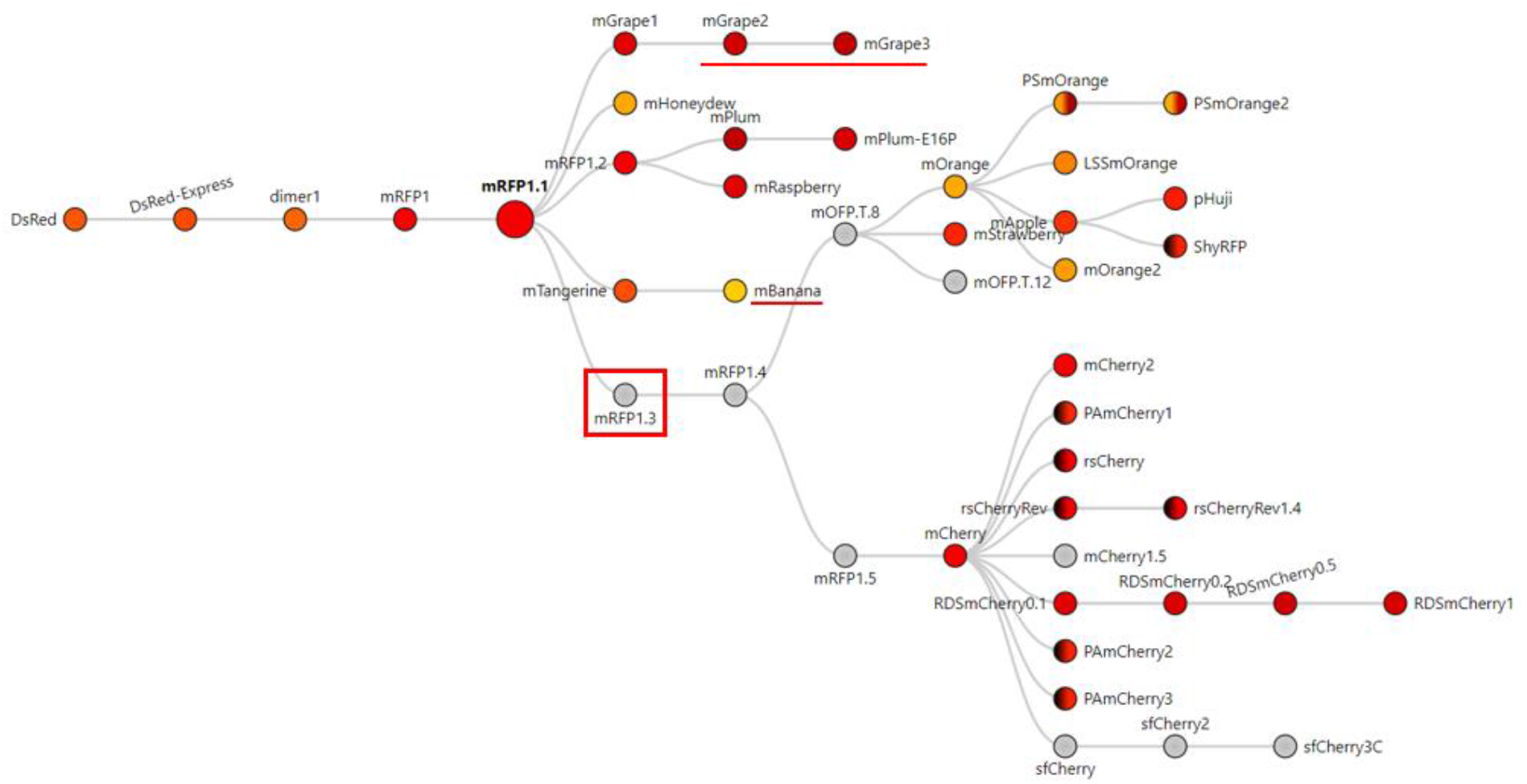
Phylogenic tree of DsRed-derived fluorescent proteins. Figure adapted from FPbase [27] with their permission. The phylogenic tree shows all the fluorescent proteins derived from mRFP1.3 that contain the problem-causing N-terminal extension. Underlined proteins are derived from mRPF1.1 and also contain the extension. Color in circles corresponds to the emission wavelength(s) of the protein. Dark hemicycle represent an off state of the protein. If colored gray, the emission maximum was not entered in the database.

**Figure 4.**
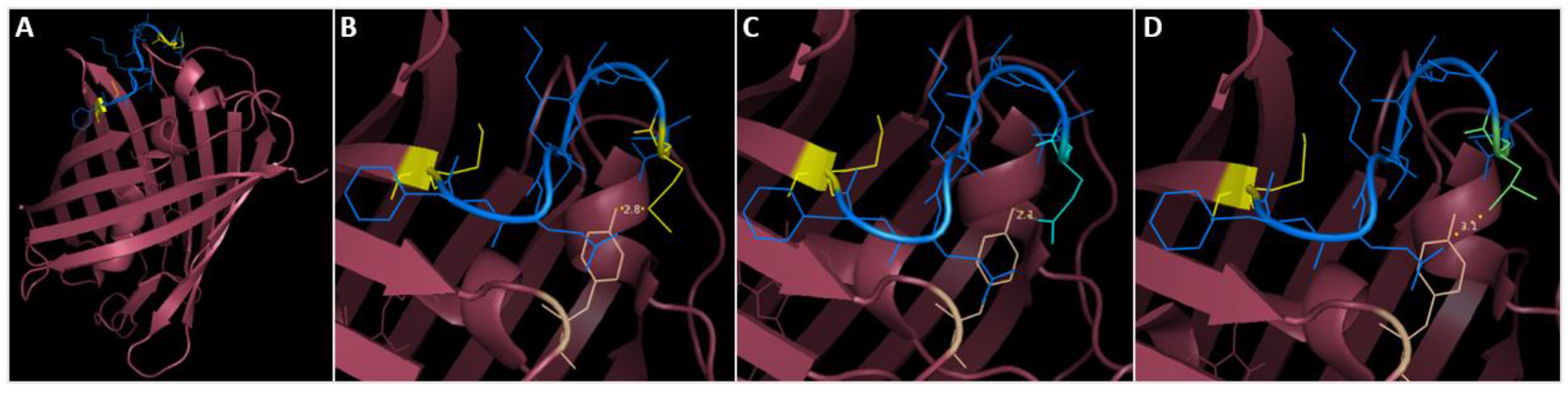
3D-structure visualization of mCherry and the substitutions provided to abolish the short isoform. (A) The segment between M10 and M17 (shown in blue, methionine in yellow) precedes the first β-sheet but provides indispensable stabilization to the protein (as mCherry V2 is not functional). The eGFP segment preceding M10 is not present on any PDB files of mCherry, potentially due to a low resolution, due to mixed isoform population. (B) M10 (yellow) stabilizes mCherry by interacting with Y43 (wheat color). (C) The M10Q substitution (in cyan) occupies a similar volume as methionine and can produce an H-bond with Y43. (D) The M10L substitution (in lime green) also mimics the steric occupancy of methionine and can interact weakly with Y43 through Van-Der-Walls interaction.

DsRed was modified into mRFP1 to overcome obligate tetramerization, improve protein maturation and modify the excitation/emission wavelength couple [16]. Consequently, mRFP1 was further modified into mRFP1.1 by the notable Q66M substitution in the chromophore to improve parameters such as photostability, quantum yield and extinction coefficient [14]. During the same round of modifications, in an attempt to improve the poor N-terminal fusion properties of mRFP1.1, the initial residues of eGFP (MVSKGEE) followed by the linker (NNMA) were added to mRFP1.1 to yield mRFP1.3 [14]. This manipulation is responsible for the apparition of the short mCherry isoform. Indeed, the linker (NNMA) provides the alternative methionine start codon and the eGFP fragment offers an SD-like sequence for ribosome entry (*Figure 1*).

The phylogenic overview of red fluorescent proteins (*Figure 3*) shows all the proteins engineered from mRFP1.3 that are affected by the dual-isoform issue. In addition to the mRFP1.3-derived proteins, mBanana, mGrape2 and mGrape3 also received the N-terminal extension described above. The problem becomes even more concerning, because it affects all commonly used variants in the red color spectrum.

### The short mCherry isoform is also expressed in other prokaryotes

In bacteria, translation initiation shares quasi-universal principles [28,29], with some variations in the SD sequences recruiting ribosomes. Since the short mCherry isoform is encoded in the *mCherry* gene sequence, it is highly likely that other prokaryotes produces the short mCherry isoform as well. In addition, depending on the codon usage of the N-terminal extension, the short *mCherry* expression may be stronger or adapted to other organisms (as demonstrated by *mCherry* with De-optimized SD (*Figure 5*). We found that the short mCherry isoform (V1) was functional in *Vibrio natriegens, Pseudomonas putida* (*Supplemental Figure S5 & Supplemental Figure S6*), and another group had found the *mCherry* ATIS occurring in *Mycoplasma* [30]. This demonstrates that the *mCherry* defect occurs across a wide range of bacteria.

**Figure 5.**
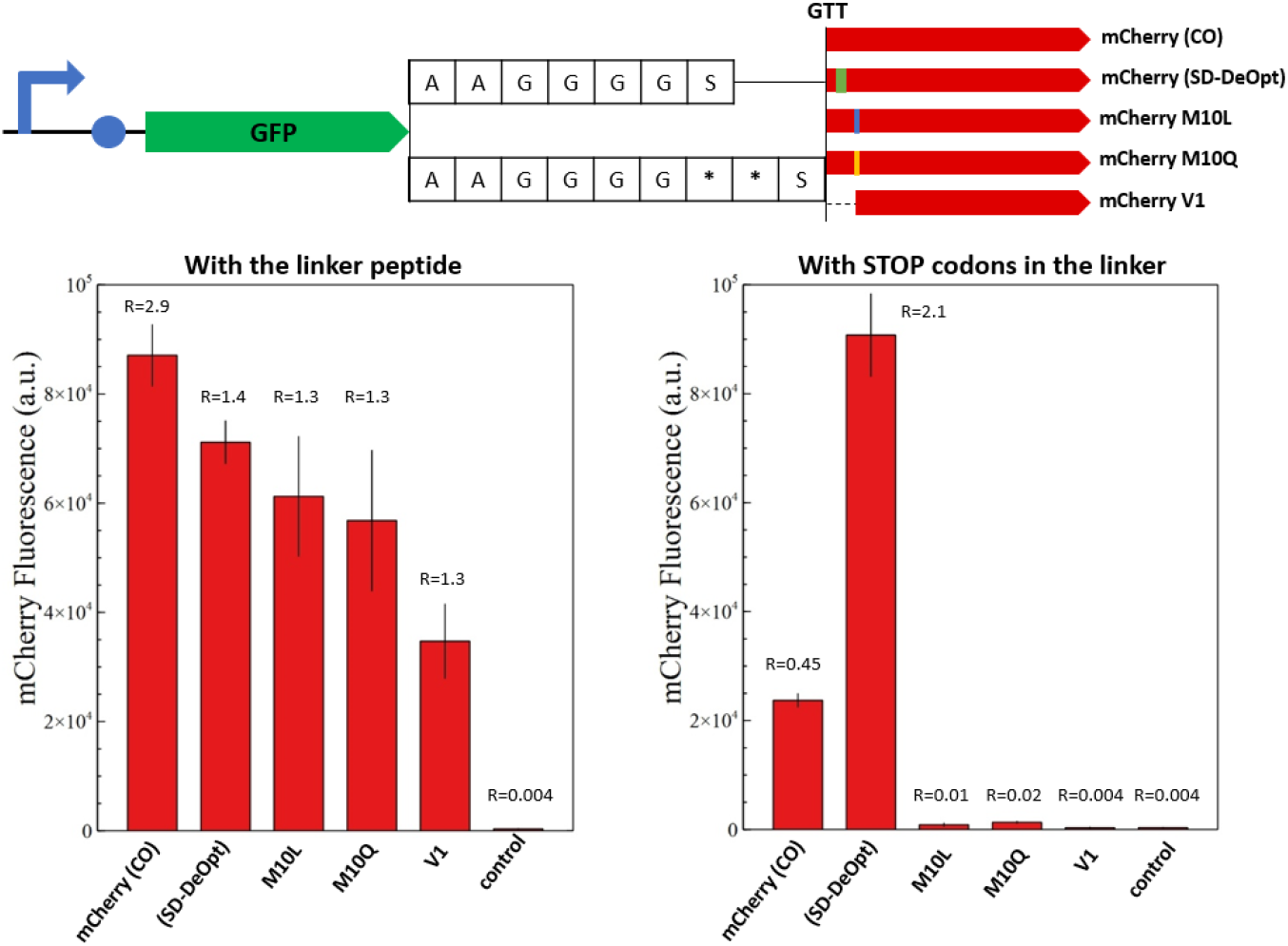
Fluorescence measurements of mCherry and its modified versions in a C-terminal fusion context. The overview of the genetic constructs is presented (top). The versions of mCherry are fused to the C-terminal end of sfGFP with a standard alanine/glycine linker or with the linker containing two stop codons. The red fluorescence of the fusion proteins with the alanine/glycine linker is presented on the left-hand side and with stop codons on the right-hand side. A plasmid expressing only *sfGFP* is used as negative control. The ratio between sfGFP and mCherry fluorescence is indicated above the corresponding histogram bar. Only mCherry (CO) and the “SD-deoptimized” version show background fluorescence with the linker containing stop codons. The M10 substitutions prove successful to abolish the short mCherry isoform in a C-terminal fusion context. Likewise, the short mCherry version V1 does not show background fluorescence but appears to be less efficient in C-terminal fusions.

### Solutions deployed to circumvent the defect in *mCherry*

In order to provide a version of *mCherry* that is usable for gene expression and protein localization without conferring background expression, we tested different solutions in a fusion protein context. The first solutions consisted in a substitution of the problem-causing M10 to glutamine or leucine, which could preserve structural properties of the protein (*Figure 5*). The second solution was simply to use *mCherry V1* as a reporter, since it resemble *mRFP1*.*1*. Fusion proteins composed of sfGFP fused with C-terminal engineered versions of mCherry were constructed to assess the performance of each solution. First, sfGFP and mCherry versions were linked with an alanine/glycine linker to estimate the C-terminal fusion properties of the engineered mCherry versions (*Figure 5*). The background expression due to the short isoform was assessed by placing two stop codons in the linker peptide right upstream the *mCherry* versions. The M10 substitutions prove successful in abolishing production of the short mCherry isoform in a C-terminal fusion context. Likewise, the short mCherry version V1 did not show any background fluorescence but appeared to be less efficient in C-terminal fusions, as reported for *mRFP1*.*1* [14].

## Conclusion

The unexpected performance of *mCherry* as a reporter in our gene expression experiments lead to the discovery of a short functional isoform of mCherry. The expression of this short isoform affects the outcome of gene expression and protein localization studies. The defect originates from a methionine residue in position 10 that is preceded by an SD-like sequence. We showed that mutation of this residue to leucine or glutamine abolishes the production of the short isoform while conserving N-terminal fusion properties.

Our findings indicate that a large proportion of the DsRed-derived proteins are affected by the production of the short isoform, due to the presence of a problem-causing linker sequence (MVSKGEE-NNMA). In addition, we identified the presence of this linker in the bright green fluorescent protein mNeonGreen [31]. We would like to alert to the distinct possibility that the production of short functional isoforms may also affect other fluorescent proteins. Indeed, the presence of methionine residues in the first 10-20 amino acids, which may act as an alternative translation start sites, is widespread among fluorescent proteins. For example, mTFP1 [32], KillerRed [33], Dronpa [34], mEosEM [35], mKelly2 [36], mGinger2 [36] and their derivatives could be affected by the production of a short isoform as observed for mCherry.

In addition, we showed that the dual-isoform issue affects various prokaryotes, as they share similar translation initiation mechanisms. Although these mechanisms differ in eukaryotes, the production of a short mCherry isoform may also occur in this group of organisms.

We demonstrate that substitution of M10 abolishes the expression of the short protein isoform. This solution should be investigated for all affected DsRed-derived proteins to ensure the accuracy and reliability as reporters. Moreover, we suggest that similar actions might be necessary for other fluorescent protein families. This work raises concerns about the outcome of studies that employed fluorescent proteins that possess an inherent background expression.

## Materials and methods

### Materials

Experiments were performed in *Escherichia coli* DH5-α (NEB), grown in LB-Lennox (Oxoid) (10 g/L casein peptone, 5 g/L yeast extract, 5 g/L NaCl with additional 15 g/L agar for plates) supplemented with 100 µg/mL ampicillin (Sigma-Aldrich). PCR amplifications were performed with Q5 polymerase (NEB). All other necessary enzymes were also purchased from NEB. Primers were ordered from Eurofins Genomics and Sigma-Aldrich. Plasmids and PCR products were purified with QIAprep plasmid Miniprep kit and QiaQuick PCR purification kit respectively (Qiagen). Plasmid Sanger sequencing was performed by Eurofins Genomics. The *E. coli* codon-optimized sequence of *mCherry* was a gift from Yanina R. Sevastsyanovich (University of Birmingham).

### Cloning and strain engineering

A pUC19 backbone containing a constitutive promoter/5’-UTR expressing *sfGFP*, generated in a previous study [18], was used as template for cloning the different versions of *mCherry*. The backbone and the different versions of *mCherry* were amplified by PCR with the respective primers presented in *Supplementary Table S1*. To test for short isoforms, *sfGFP* was replaced by the codon-optimized *mCherry* gene or its shorter versions (V1, V2 and V3) by Golden gate assembly [37]. To build fusion constructs, the different *mCherry* versions were fused downstream *sfGFP* using Golden Gate assembly.

The Golden Gate assembly mixture was chemically transformed into competent *E. coli* by heat shock (45s at 42°C). Cells were plated on LB plates containing 100 µg/mL ampicillin and grown overnight at 37°C. Positive clones were grown in 5 mL LB supplemented with 100 µg/mL ampicillin and their respective plasmids were purified. DNA sequences were confirmed by Sanger sequencing.

### Fluorescence measurements

Each *E. coli* strain bearing a given plasmid was inoculated into LB supplemented with 100 µg/mL ampicillin on 96-well plates and incubated overnight at 37°C with 800 rpm agitation in a Multitron Pro plate-shaking incubator (Infors HT). Fluorescence was measured on an Infinite M200 Pro TECAN fluorimeter (Noax Lab AS). The excitation/emission wavelengths were 488/525 nm for GFP (gain 67) and 576/610 nm for mCherry (gain 97). Fluorescence intensity was normalized by OD _600_ of the corresponding well.

### Bioinformatics analysis

The amino acid sequence of mCherry-(CO) was used as template to perform preliminary protein t BLAST searches (blast.ncbi.nlm.nih.gov/Blast.cgi). Phylogeny of red-fluorescent proteins was consulted on FPbase [27] (fpbase.org).

### Proteomics analysis

The strain carrying the constitutively expressed *mCherry V1* was grown overnight in 50 mL of LB media supplemented with 100 µg/mL ampicillin at 37°C-225 rpm. Cells were harvested by centrifugation (5000 rpm, 5 min) and resuspended in lysis buffer (50 mM Tris-HCl, 50 mM NaCl, 0.05 % Triton X-100, pH 8.0) supplemented with 1 tablet EDTA-free cOmplete ULTRA protease inhibitor (Roche). Cell debris were eliminated by centrifugation (7500 rpm, 15 min) and the soluble fraction was collected.

### Sample Preparation for LC-MS – Protein Digestion

2 samples of 200 µl cell lysate each were separated for LC-MS analysis, one to be digested by Trypsin the other by both Trypsin and Lys-C. Soluble proteins were precipitated by methanol/chloroform/water precipitation. Briefly, 800 µl methanol was added to the sample, followed by the addition of 200µl chloroform and vortexing. After addition of 600 µl ultra pure water (18 MΩ), samples were thoroughly mixed by vortexing and centrifuged at 16000g for 2 minutes. The upper layer was discarded without disturbing the protein layer and further 800 µl methanol was added, followed by vortexing and centrifugation as above. After removing the supernatant, the protein pellet was air dried for 10 minutes, it was then reconstituted in 150 µl of 50 mM ammonium bicarbonate (BioUltra; Sigma-Aldrich, Germany). Next, proteins were reduced with 1.5 µmol of DTT (Sigma-Aldrich, Canada) for 20 minutes at 70 degrees Celsius, then they were brought back to room temperature and then they were alkylated using 6 µmoles of iodoacetamide (BioUltra; Sigma-Aldrich, USA) in the dark at room temperature for 30 minutes. Excess iodoacetamide was quenched by adding 3.5 µmoles of DTT (Sigma-Aldrich, Canada) and incubating in the dark for 20 min at room temperature. Finally, proteins were digested by endoproteinase at 37 °C overnight, one sample digested by 1.25 µg trypsin and the other by 1.25 µg trypsin and 1.25 µg of Lys-C. After overnight digestion, 5 µl of formic acid was added to each sample and then the peptides were dried in a vacuum concentrator at 60 degrees Celsius.

### Sample Preparation for LC-MS – Peptide Desalting

Samples were resuspended in 100 µl 0.1 % formic acid and desalted in C18 stage tip columns, unless otherwise specified chemicals are Optima grade from Fisher Chemicals and centrifugations are at 2000g. Briefly, Stage tip columns consisting of three C18 plugs (Empore C18 47mm SPE Disks, 3M, USA) were made and activated with 50 µL methanol by centrifugation for 2 minutes, the methanol activation was repeated. Then the Stage tip was equilibrated with 60 µL of 0.1% formic acid in water by centrifugation for 2 minutes, the equilibration was repeated. Peptide-samples were centrifuged (16000g for 25 minutes), supernatants were loaded onto stage tip columns and centrifuged for 4 minutes, flow-through solutions were reloaded to stage tip columns. Stage tip columns were washed with 60 µL 0.1% formic acid by centrifugation for 2 minutes, the wash was repeated. Peptides were then eluted from the stage tip column using 40 µL 70% acetonitrile, 0.1% formic acid, centrifugation for 2 minutes, the elution step was repeated. Finally, desalted peptides were dried in a vacuum concentrator at 60 degrees Celsius and stored at -20 degrees Celsius until LC-MS analysis.

### LC-MS analysis

Dried peptides were reconstituted in 50 µL 0.1% formic acid in water and shaken at 6 degrees Celsius at 900 rpm for 1.5 hour. Samples were centrifuged at 16000g for 10 minutes and 40 µL supernatants were transferred to MS-vials for LC-MS analysis. LC-MS analysis was performed on an EASY-nLC 1200 UPLC system (Thermo Scientific) interfaced with an Q Exactive mass spectrometer (Thermo Scientific) via a Nanospray Flex ion source (Thermo Scientific). Peptides were injected onto an Acclaim PepMap100 C18 trap column (75 μm i.d., 2 cm long, 3 μm, 100 Å, Thermo Scientific) and further separated on an Acclaim PepMap100 C18 analytical column (75 μm i.d., 50 cm long, 2 μm, 100 Å, Thermo Scientific) using a 180-minute multi-step gradient (150 min 2%-40% B, 15 min 40%-100% B, 15 min at 100% B; where B is 0.1 % formic acid and 80% CH_3_CN and A is 0.1 % formic acid) at 250 nl/min flow. Peptides were analyzed in positive ion mode under data dependent acquisition using the following parameters: Electrospray voltage 1.9 kV, HCD fragmentation with normalized collision energy 28. Each MS scan (200 to 2000 m/z, 2 m/z isolation width, profile) was acquired at a resolution of 70,000 FWHM in the Orbitrap analyzer, followed by MS/MS scans at resolution 17,500 (2 m/z isolation width, profile) triggered for the 12 most intense ions, with a 30 s dynamic exclusion and analyzed in the Orbitrap analyzer. Charge exclusion was set to unassigned, 1, >4.

### Processing of LC-MS Data

Proteins were identified by processing the LC-MS data using Thermo Scientific Proteome Discoverer (Thermo Scientific) version 2.5. The following search parameters were used: enzyme specified as trypsin with maximum two missed cleavages allowed; acetylation of protein N-terminus with methionine loss, oxidation of methionine, and deamidation of asparagine/glutamine were considered as dynamic and carbamidomethylation of cysteine as static post-translational modifications; precursor mass tolerance of 10 parts per million with a fragment mass tolerance of 0.02 Da. Sequest HT node queried the raw files against sequences for mCherry (original and short), E. coli (strain K-12) proteins downloaded from Uniprot (https://www.uniprot.org/proteomes/UP000000625) in September 2020 and a common LC-MS contaminants database. For downstream analysis of these peptide-spectrum matches (PSMs), for protein and peptide identifications the PSM FDR was set to 1% and as high and 5% as medium confidence, thus only unique peptides with these confidence thresholds were used for final protein group identification and to label the level of confidence, respectively.

### Protein 3D structure modelling

The 3D structure modelling of mCherry was done with the software Pymol and the PDB file 2H5Q. In all mCherry PDB files (2H5Q, 6YLM, 6IR1&2, 6MZ3, 4ZIN), the first eight amino acids are unmodeled, residue 9 and 10 are computationally added but M10 is mis-modeled. We corrected residue 10 to model methionine. Then we changed M10 into glutamine and leucine on the 3D structure with Pymol. Rotamers with 2-3Å proximity with Y43 are shown to model their interactions.

## Supplemental Figures and Tables

**Supplemental Table S1.**
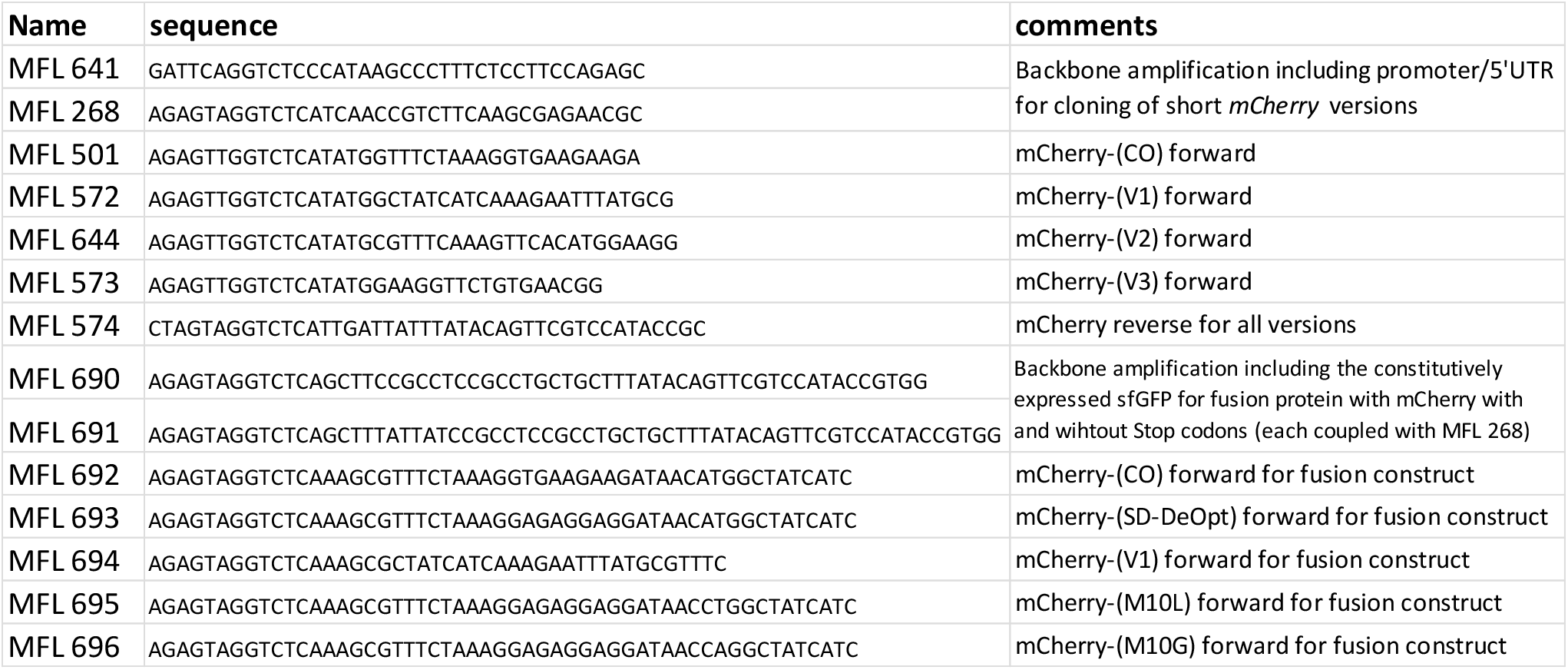
Primer list.

**Supplemental Figure S1.**
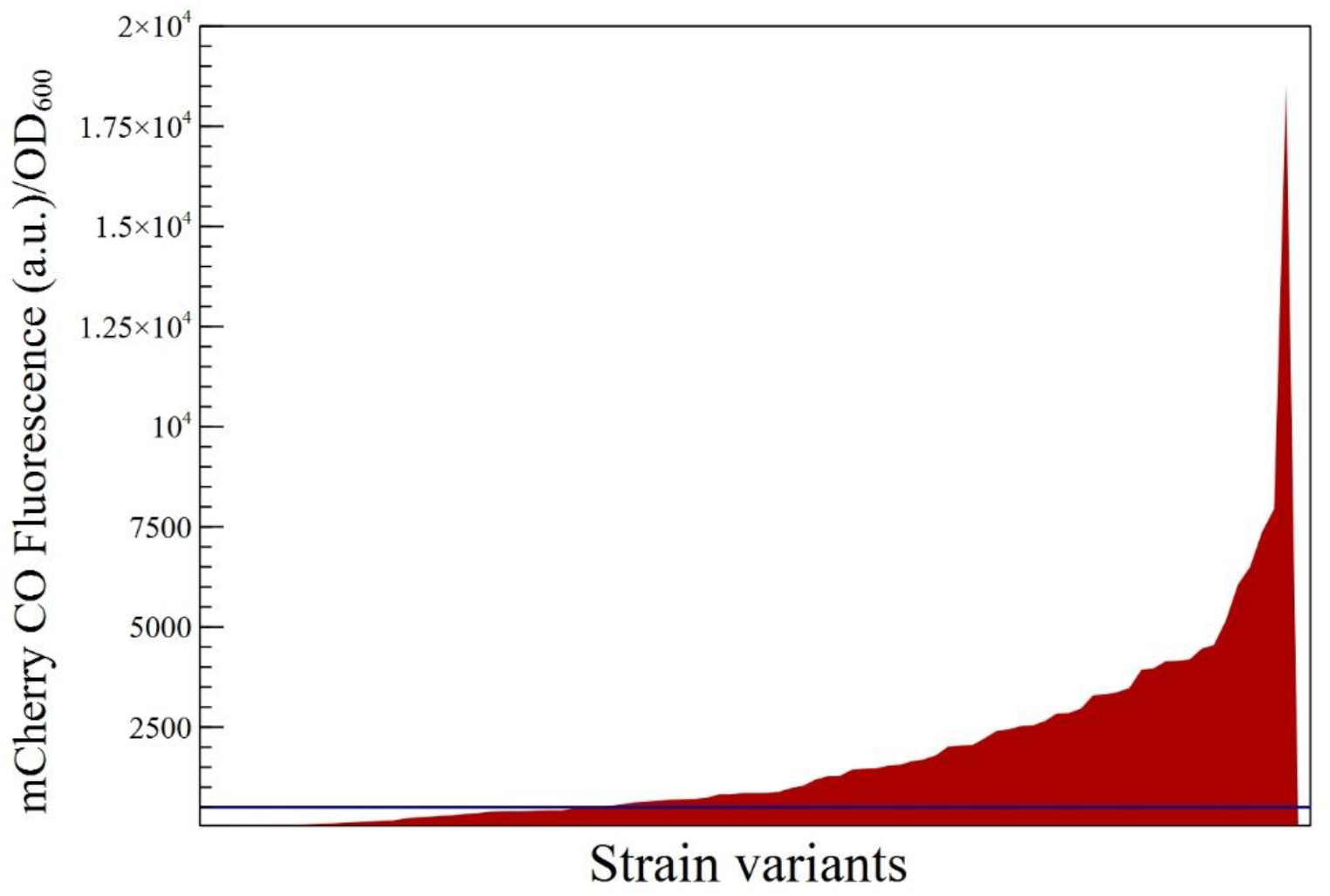
Quantification of fluorescent *E. coli* strains carrying the 200N random library in front of the original *mCherry* gene. The 200N library was cloned in front of the original mCherry (CO) and random clones were grown for fluorescence measurement. The number of positives clones obtained with *mCherry* exceeds significantly the usual 30-40% efficiency of the method. Here, the significance threshold (blue line) indicates that 65% of clones express mCherry.

**Supplemental Figure S2.**
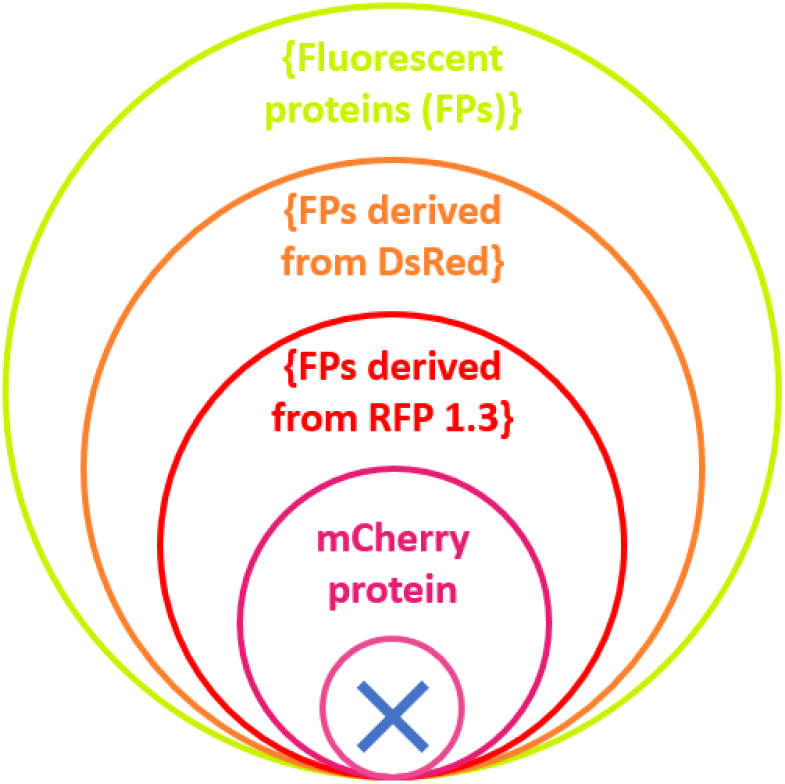
Representation of Russell’s paradox applied to mCherry. Each circle represents a set of fluorescent proteins that contains a smaller set. The mCherry protein is supposedly the smallest set. However, the circle with a cross represents Russell’s paradox in the form of a short mCherry isoform contained in itself.

**Supplemental Figure S3.**
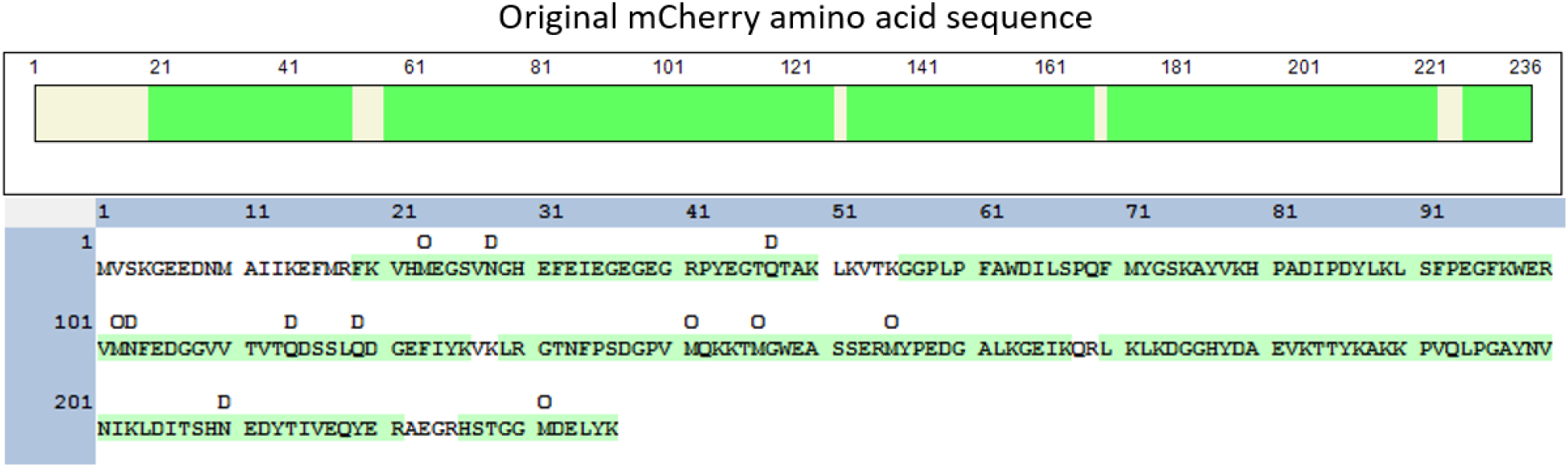
Peptide coverage of the mCherry V1 fluorescent protein found by proteomics. Peptides identified by LC-MS analysis are displayed in green. The main initial fragment from G5 to K14 is absent, which confirms that the fluorescent isoform of mCherry lacks the N-terminal region. The fragment between M10 and M17 (difference between mCherry V1 and V2) was not detected due to the the cleavage site K14. However, mCherry V2 did not show fluorescence, thus mCherry V1 is the only short functional isoform.

**Supplemental Figure S4.**
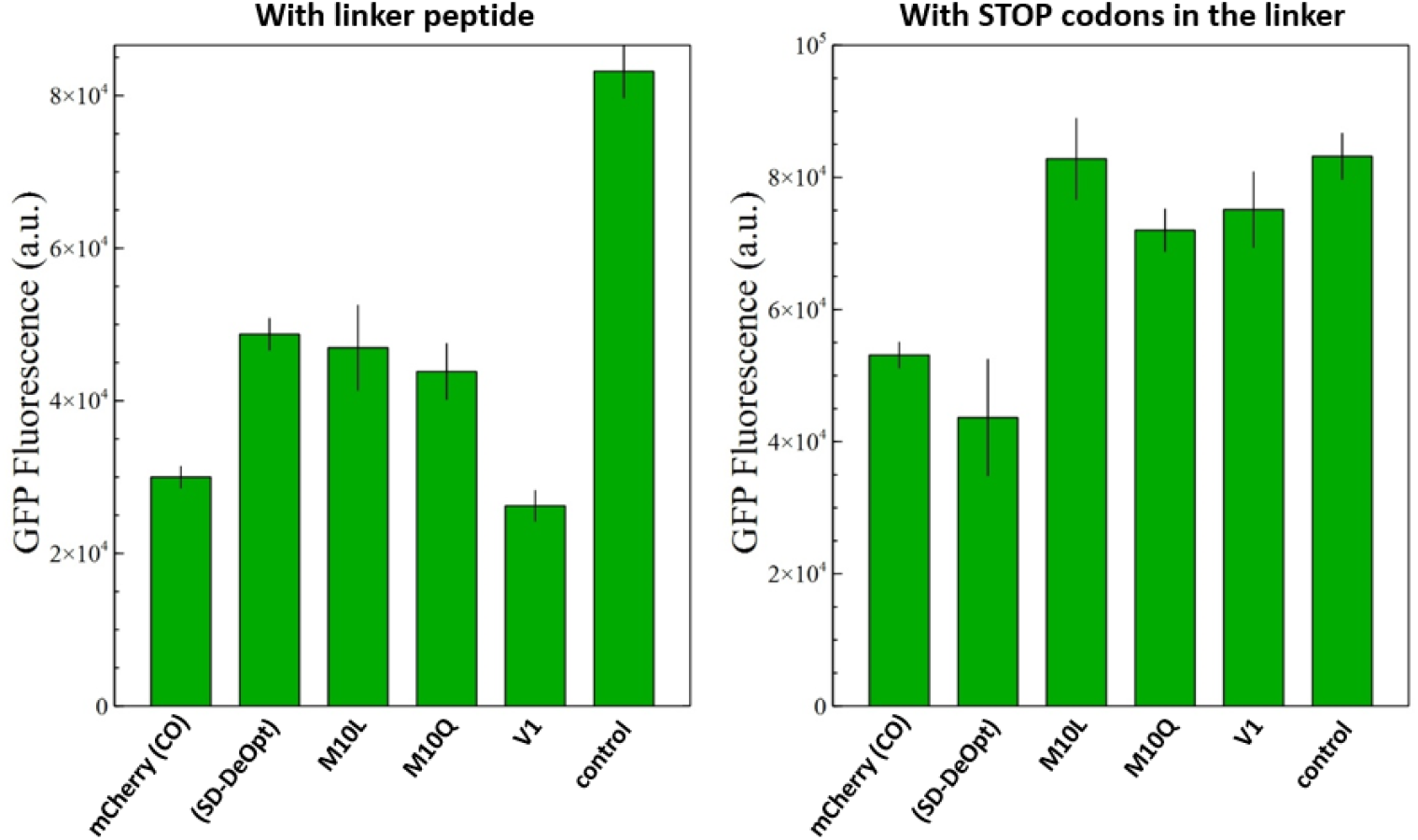
Fluorescence measurements of sfGFP from the fusions with the different versions of mCherry. The sfGFP fluorescence was used to normalize the mCherry fluorescence and provide a relative ratio for comparison between mCherry versions (in *Figure 5*). A plasmid expressing only sfGFP is used as negative control.

**Supplemental Figure S5.**
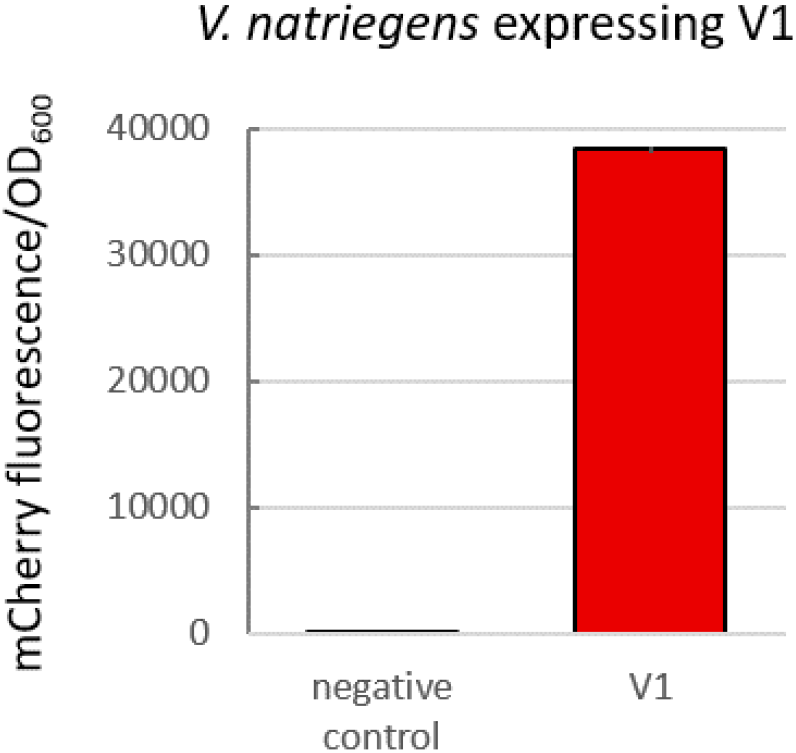
Fluorescence measurement of the short mCherry isoform V1 in *V. natriegenes*. The short *mCherry* version was cloned onto a pACYC backbone and constitutively expressed in *V. natriegens*. The short gene produced a functional protein detected by fluorescence.

**Supplemental Figure S6.**
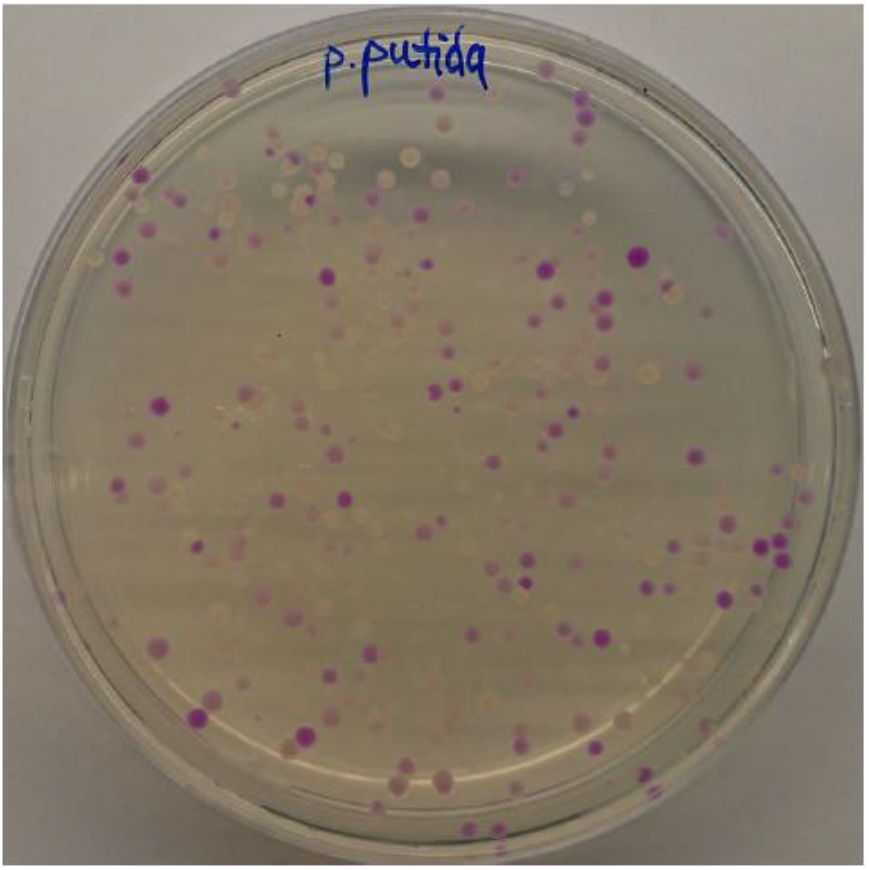
Photograph of *P. putida* expressing *mCherry* original gene with the 200N random DNA library. The percentage of positive clones exceeds significantly the usual efficiency of the method, suggesting that the ATIS is also active in *P. putida*.

## Notes

### Competing Interest Statement

The authors have declared no competing interest.

## References

[1] J. van Heijenoort, From Frege to Gödel: a source book in mathematical logic, 1879–1931, Harvard University Press, 2002.

[2] D. Jaquette, Personal Tragedy and a Philosophical Hiatus (1904–1917), in: Frege A Philos. Biogr., Cambridge University Press, 2019. https://doi.org/10.1017/9781139033725.015.

[3] D.M. Chudakov, M. V. Matz, S. Lukyanov, K.A. Lukyanov, Fluorescent Proteins and Their Applications in Imaging Living Cells and Tissues, Physiol. Rev. 90 (2010). https://doi.org/10.1152/physrev.00038.2009.

[4] R.N. Day, M.W. Davidson, The fluorescent protein palette: tools for cellular imaging, Chem. Soc. Rev. 38 (2009). https://doi.org/10.1039/b901966a.

[5] R. Ranganathan, M.A. Wall, M. Socolich, The structural basis for red fluorescence in the tetrameric GFP homolog DsRed, Nat. Struct. Biol. 7 (2000). https://doi.org/10.1038/81992.

[6] R.M. Wachter, J.L. Watkins, H. Kim, Mechanistic Diversity of Red Fluorescence Acquisition by GFP-like Proteins, Biochemistry. 49 (2010). https://doi.org/10.1021/bi100901h.

[7] L.A. Gross, G.S. Baird, R.C. Hoffman, K.K. Baldridge, R.Y. Tsien, The structure of the chromophore within DsRed, a red fluorescent protein from coral, Proc. Natl. Acad. Sci. 97 (2000) 11990–11995. http://www.pnas.org.

[8] X. Shu, N.C. Shaner, C.A. Yarbrough, R.Y. Tsien, S.J. Remington, Novel Chromophores and Buried Charges Control Color in mFruits, Biochemistry. 45 (2006). https://doi.org/10.1021/bi060773l.

[9] M. V. Matz, A.F. Fradkov, Y.A. Labas, A.P. Savitsky, A.G. Zaraisky, M.L. Markelov, S.A. Lukyanov, Fluorescent proteins from nonbioluminescent Anthozoa species, Nat. Biotechnol. 17 (1999). https://doi.org/10.1038/13657.

[10] F. V. Subach, V. V. Verkhusha, Chromophore Transformations in Red Fluorescent Proteins, Chem. Rev. 112 (2012). https://doi.org/10.1021/cr2001965.

[11] A.S. Mishin, F. V. Subach, I. V. Yampolsky, W. King, K.A. Lukyanov, V. V. Verkhusha, The first mutant of the Aequorea victoria green fluorescent protein that forms a red chromophore, Biochemistry. 47 (2008) 4666–4673. https://doi.org/10.1021/bi702130s.

[12] Y.A. Labas, N.G. Gurskaya, Y.G. Yanushevich, A.F. Fradkov, K.A. Lukyanov, S.A. Lukyanov, M. V. Matz, Diversity and evolution of the green fluorescent protein family, Proc. Natl. Acad. Sci. 99 (2002). https://doi.org/10.1073/pnas.062552299.

[13] B.J. Bevis, B.S. Glick, Rapidly maturing variants of the Discosoma red fluorescent protein (DsRed), (2002). http://biotech.nature.com.

[14] N.C. Shaner, R.E. Campbell, P.A. Steinbach, B.N.G. Giepmans, A.E. Palmer, R.Y. Tsien, Improved monomeric red, orange and yellow fluorescent proteins derived from Discosoma sp. red fluorescent protein, Nat. Biotechnol. 22 (2004) 1567–1572. https://doi.org/10.1038/nbt1037.

[15] E.A. Rodriguez, R.E. Campbell, J.Y. Lin, M.Z. Lin, A. Miyawaki, A.E. Palmer, X. Shu, J. Zhang, R.Y. Tsien, The Growing and Glowing Toolbox of Fluorescent and Photoactive Proteins, Trends Biochem. Sci. 42 (2017) 111–129. https://doi.org/10.1016/j.tibs.2016.09.010.

[16] R.E. Campbell, O. Tour, A.E. Palmer, P.A. Steinbach, G.S. Baird, D.A. Zacharias, R.Y. Tsien, A monomeric red fluorescent protein, Proc. Natl. Acad. Sci. 99 (2002). https://doi.org/10.1073/pnas.082243699.

[17] M. Lauff, R. Hofer, Proteolytic enzymes in fish development and the importance of dietary enzymes, Aquaculture. (1984). https://doi.org/10.1016/0044-8486(84)90298-9.

[18] R. Lale, L. Tietze, J. Nesje, I. Onsager, K. Engelhardt, C. Fai Alex, M. Akan, N. Hummel, J. Kalinowski, C. Rückert, M. Frank, A universal method for gene expression engineering, (n.d.). https://doi.org/10.1101/644989.

[19] C. Fritsch, A. Herrmann, M. Nothnagel, K. Szafranski, K. Huse, F. Schumann, S. Schreiber, M. Platzer, M. Krawczak, J. Hampe, M. Brosch, Genome-wide search for novel human uORFs and N-terminal protein extensions using ribosomal footprinting, Genome Res. 22 (2012). https://doi.org/10.1101/gr.139568.112.

[20] J. Wan, S.-B. Qian, TISdb: a database for alternative translation initiation in mammalian cells, Nucleic Acids Res. 42 (2014). https://doi.org/10.1093/nar/gkt1085.

[21] J.L. Wegrzyn, T.M. Drudge, F. Valafar, V. Hook, Bioinformatic analyses of mammalian 5’-UTR sequence properties of mRNAs predicts alternative translation initiation sites, BMC Bioinformatics. 9 (2008). https://doi.org/10.1186/1471-2105-9-232.

[22] K. Nakahigashi, Y. Takai, M. Kimura, N. Abe, T. Nakayashiki, Y. Shiwa, H. Yoshikawa, B.L. Wanner, Y. Ishihama, H. Mori, Comprehensive identification of translation start sites by tetracycline-inhibited ribosome profiling, DNA Res. 23 (2016) 193–201. https://doi.org/10.1093/dnares/dsw008.

[23] P. Trulley, G. Snieckute, D. Bekker-Jensen, M.B. Menon, R. Freund, A. Kotlyarov, J. V. Olsen, M.D. Diaz-Muñoz, M. Turner, S. Bekker-Jensen, M. Gaestel, C. Tiedje, Alternative Translation Initiation Generates a Functionally Distinct Isoform of the Stress-Activated Protein Kinase MK2, Cell Rep. 27 (2019) 2859–2870.e6. https://doi.org/10.1016/j.celrep.2019.05.024.

[24] A.J. Ozin, T. Costa, A.O. Henriques, J. Moran, Alternative translation initiation produces a short form of a spore coat protein in Bacillus subtilis, J. Bacteriol. 183 (2001) 2032–2040. https://doi.org/10.1128/JB.183.6.2032-2040.2001.

[25] S.C. Bernier, L.P. Morency, R. Najmanovich, C. Salesse, Identification of an alternative translation initiation site in the sequence of the commonly used Glutathione S-Transferase tag, J. Biotechnol. 286 (2018) 14–16. https://doi.org/10.1016/j.jbiotec.2018.09.003.

[26] S.M. Chabregas, D.D. Luche, M.A. Van Sluys, C.F.M. Menck, M.C. Silva-Filho, Differential usage of two in-frame translational start codons regulates subcellular localization of Arabidopsis thaliana THI1, J. Cell Sci. 116 (2003) 285–291. https://doi.org/10.1242/jcs.00228.

[27] T.J. Lambert, FPbase: a community-editable fluorescent protein database, Nat. Methods. 16 (2019) 277–278. https://doi.org/10.1038/s41592-019-0352-8.

[28] M. V. Rodnina, Translation in prokaryotes, Cold Spring Harb. Perspect. Biol. 10 (2018). https://doi.org/10.1101/cshperspect.a032664.

[29] S. Nakagawa, Y. Niimura, K.I. Miura, T. Gojobori, Dynamic evolution of translation initiation mechanisms in prokaryotes, Proc. Natl. Acad. Sci. U. S. A. 107 (2010) 6382–6387. https://doi.org/10.1073/pnas.1002036107.

[30] P. Carroll, J. Muwanguzi-Karugaba, E. Melief, M. Files, T. Parish, Identification of the translational start site of codon-optimized mCherry in Mycobacterium tuberculosis, BMC Res. Notes. 7 (2014). https://doi.org/10.1186/1756-0500-7-366.

[31] N.C. Shaner, G.G. Lambert, A. Chammas, Y. Ni, P.J. Cranfill, M.A. Baird, B.R. Sell, J.R. Allen, R.N. Day, M. Israelsson, M.W. Davidson, J. Wang, A bright monomeric green fluorescent protein derived from Branchiostoma lanceolatum, Nat. Methods. 10 (2013). https://doi.org/10.1038/nmeth.2413.

[32] H. Ai, J.N. Henderson, S.J. Remington, R.E. Campbell, Directed evolution of a monomeric, bright and photostable version of Clavularia cyan fluorescent protein: structural characterization and applications in fluorescence imaging, Biochem. J. 400 (2006). https://doi.org/10.1042/BJ20060874.

[33] M.E. Bulina, D.M. Chudakov, O. V Britanova, Y.G. Yanushevich, D.B. Staroverov, T. V Chepurnykh, E.M. Merzlyak, M.A. Shkrob, S. Lukyanov, K.A. Lukyanov, A genetically encoded photosensitizer, Nat. Biotechnol. 24 (2006). https://doi.org/10.1038/nbt1175.

[34] R. Ando, Regulated Fast Nucleocytoplasmic Shuttling Observed by Reversible Protein Highlighting, Science (80-.). 306 (2004). https://doi.org/10.1126/science.1102506.

[35] Z. Fu, D. Peng, M. Zhang, F. Xue, R. Zhang, W. He, T. Xu, P. Xu, mEosEM withstands osmium staining and Epon embedding for super-resolution CLEM, Nat. Methods. 17 (2020). https://doi.org/10.1038/s41592-019-0613-6.

[36] T.M. Wannier, S.K. Gillespie, N. Hutchins, R.S. McIsaac, S.-Y. Wu, Y. Shen, R.E. Campbell, K.S. Brown, S.L. Mayo, Monomerization of far-red fluorescent proteins, Proc. Natl. Acad. Sci. 115 (2018). https://doi.org/10.1073/pnas.1807449115.

[37] C. Engler, R. Kandzia, S. Marillonnet, A one pot, one step, precision cloning method with high throughput capability, PLoS One. 3 (2008). https://doi.org/10.1371/journal.pone.0003647.

